# Complementary Performances of Convolutional and Capsule Neural Networks on Classifying Microfluidic Images of Dividing Yeast Cells

**DOI:** 10.1101/852566

**Authors:** Mehran Ghafari, Justin Clark, Hao-Bo Guo, Ruofan Yu, Yu Sun, Weiwei Dang, Hong Qin

## Abstract

Microfluidic-based assays have become effective high-throughput approaches to examining replicative aging of budding yeast cells. Deep learning may offer an efficient way to analyze a large number of images collected from microfluidic experiments. Here, we compare three deep learning architectures to classify microfluidic time-lapsed images of dividing yeast cells into categories that represent different stages in the yeast replicative aging process. We found that convolutional neural networks outperformed capsule networks in terms of accuracy, precision, and recall. The capsule networks had the most robust performance at detecting one specific category of cell images. An ensemble of three best-fitted single-architecture models achieves the highest overall accuracy, precision, and recall due to complementary performances. In addition, extending classification classes and augmentation of the training dataset can improve the predictions of the biological categories in our study. This work lays a useful framework for sophisticated deep-learning processing of microfluidics-based assays of yeast replicative aging.

## Introduction

The budding yeast *Saccharomyces cerevisiae* is an effective model for studying cellular aging [1, 2]. The replicative lifespan of a yeast mother cell is defined as the total number of cell divisions accomplished, or the number of daughter cells produced throughout its lifetime. Microfluidics is a fast-growing tool for high-throughput applications in chemical, biological, optics, and information technology, including single-cell imaging analysis [3]. Typically, microfluidic images are taken in time intervals with relatively low resolution compared to confocal microscopic images that are often of high resolution, rendering unique challenges for microfluidic imaging [4]. Capturing the full progression of cellular replicative lifespans requires identifying both mother cells and daughter cells in full cell cycles. However, the full automation of this process is often hindered by low image resolution, demanding time-consuming, manual classifications of yeast replicative lifespans [5].

Deep learning as a sub-field of machine learning has been applied in a wide range of applications, and its developments are mostly driven by both computational capacity and the accessibility of datasets. The development of deep learning is driven by its ability to understand and infer information from data such as speech, text, and images [6]. Medical and healthcare application areas, especially the medical images that play key roles in health diagnosis, may also benefit from machine learning detection and classification [7]. In recent years, deep learning has increased in efficacy for image classification and is now a popular method for parsing image information [8]. Many innovations have been driven by creating models that perform well on benchmark datasets such as MNIST (60,000 handwritten digits for training in a 28×28-dimensional vector space), CIFAR10 (60,000 commonly used images in a 32×32-dimensional vector space), CIFAR100 (500 training images grouped into 20 classes), and ImageNet (100,000+ phrases and around 1,000 images for each phrase) [9]. The basic idea of deep learning is to create or “learn” a function that can map a high-dimensional input space into an output vector.

In classification, the size of the output vector depends on the number of classes, while regression typically has a scalar output. In image classification problems, the convolutional neural network (CNN) is the primary type of deep learning model employed. A variety of CNN approaches have been proven useful for image classification, because they are designed for 2-dimensional (or higher) input tensors [10]. In addition, the proximity of pixels in the input images is taken into consideration, which helps CNNs learn how pixels are oriented relative to each other. One of the major drawbacks of CNNs is that they require a large amount of training samples, which is rooted in the architectural designs of CNNs [11].

A fundamentally different type of deep learning architecture, named CapsNet, was proposed to learn from fewer training samples than its traditional CNN counterparts. The recently proposed CapsNet architecture [12] is known as capsule networks with dynamic routing. The model is promising in image classification applications in datasets with limited data [13]. The success of the model lies in its ability to preserve additional information from input images by utilizing convolutional strides and dynamic routing instead of a max pooling layer. A recent study showed that CapsNet could classify fluorescent microscopic images [31].

Our work here focuses on comparing deep-learning classification methods of microfluidics images of dividing yeast cells. We compare three deep-learning neural network approaches, including CapsNet, to classify microfluidic trap images into four biological categories. The main purpose of this work is to develop a method to accurately classify microfluidic images from a small and noisy dataset. Due to data limitations, we trained each model with consideration of the effect of data augmentation. Finally, we showed that an ensemble of the top three models performs better than using each individual model alone, leading to a good “collaboration” among these models. In addition, data augmentation and splitting a class into two classes could be an effective approach for some models based on the type of dataset and model architecture.

## Materials and methods

### Hardware and hyperparameters

The models were trained and tested on NVIDIA Tesla P100 GPU. We performed a basic grid search on six hyper-parameters: (1) the number of routing iterations, (2) learning rate, (3) batch size, (4) whether to add noise to training images, (5) the number of epochs in training, and (6) whether augmentation was applied or not. The options of the hyper-parameter grid search are listed in S1 Table of the supporting information (SI). In general, a total of 108 combinations were initially tested.

### Dataset

The dataset is collected from a recent version of high-throughput yeast aging analysis (HYAA) chips experimental work [14]. Each time-lapsed image has a resolution of 1280×960 and contains approximately 114 traps as shown in Fig 1 (a). In general, traps are designed to hold a single dividing mother cell. The inlet width, outlet width, and height of each trap are 6, 3, and 5 micrometers, respectively. The outlet is wide enough to allow smaller daughter cells to slip through the trap outlet but narrow enough to withhold the bigger mother cell. Due to cell migrations (see S1 Fig (a)), low resolution, image intensity variations (see S1 Fig (b)), and difficulties in alignment, each time-lapsed image is partitioned into sub-images of 60×60 pixels, for an individual trap with respect to the boundary of its neighbor-traps as shown in Fig 1 (b). After partitioning, any individual trap typically contains 391 time-lapsed images with 10-minute intervals, which is illustrated in Fig 1 (c).

**Fig 1.**
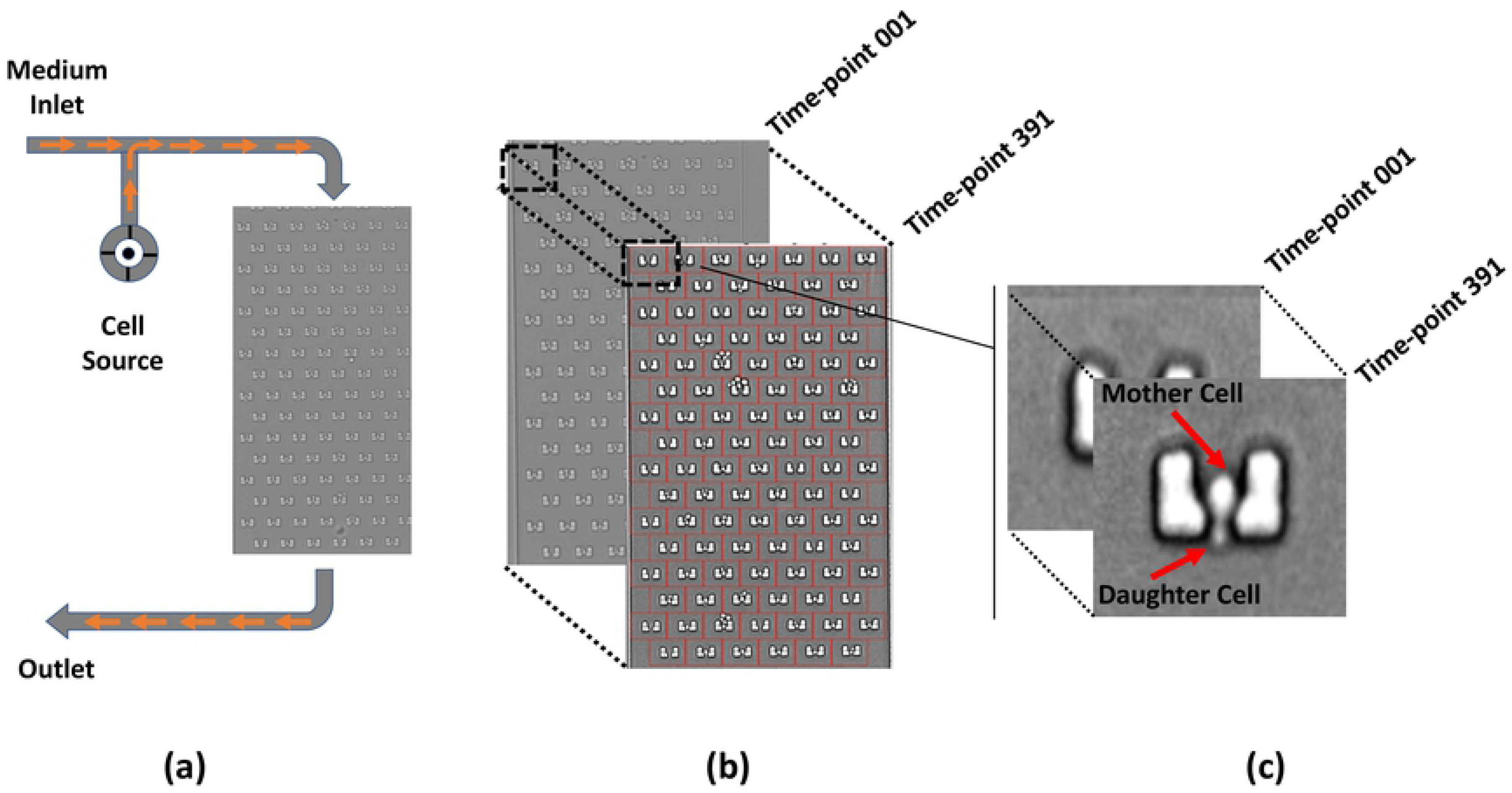
The Architecture of a Microfluidic Device. (a) Single-channel microfluidic device with medium flow direction. Cells are inserted from cell source and joint medium before reaching microfluidic traps. (b) Partitioning 114 traps of each microfluidic time-lapsed images. (c) Time-lapsed cropped images of a single trap in dimension of 60×60 pixels.

We initially categorized images based on the number of cells available at each trap and cell position. All class categories are labeled as follows: a trap with no cell (nC), a trap with a single mother cell (mC), a trap with mother and upward-oriented daughter cells (mduC), a trap with mother and downward-oriented daughter cells (mddC), and a trap with more than two cells (exC). We called all of these categories “the 5 computed classes,” as illustrated in Fig 2 (a). The exC class is necessary because it is difficult to determine the appearance of extra cells without knowledge from its immediate neighbor images; as a consequence, a trap with more than two cells will be ignored in the classification process. However, for easier understanding from a biological point of view, mddC and mduC classes are merged and labeled mdC after the testing process. In other words, all training and testing datasets are based on 5 classes and results are displayed as only 4 classes. Since mddC and mduC classes are merged together, we called these new categories “the 4 biological classes,” which include nC, mC, mdC, and exC as shown in Fig 2 (b). Examples of mddC and mduC classes with indication of cell position are shown in Fig 2 (c).

**Fig 2.**
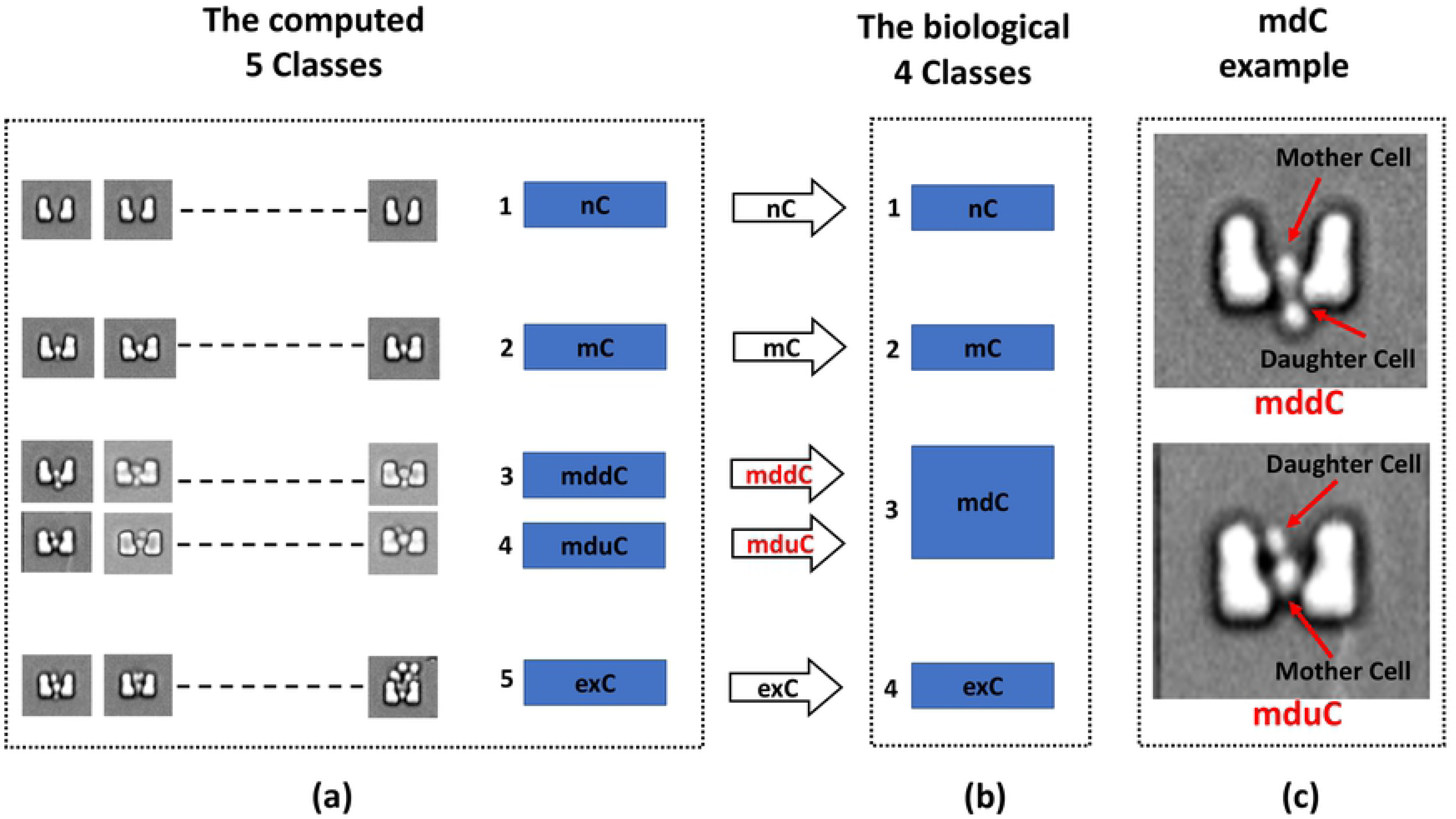
Class categories with indication of results labeling for each class. (a) 5 computed categories including nC, mC, mddC, mduC, and exC classes. (b) 4 biological categories including nC, mC, mdC, and exC classes. (c) An example of mddC and mduC for daughter cell orientation around a trap-center mother cell.

### A 2-layered architecture, CNN-2

The two-layered architecture CNN that has two convolution layers represents one of the most simplified CNN models, and it is also termed as the baseline CNN architecture. We chose this model for its simplicity, and we refer to it as the CNN-2 in the present work. The kernel size is 3×3 and batch normalization is applied to the both layers. The stride for the second layer is 2, and the activation function is ReLU for this model. Moreover, the input image size is 60×60 pixels and no image enhancement method is applied. A 2×2 kernel size used for max-pooling and 25% dropout applied for the second layer as the model architecture is shown in Fig 3 (a). We trained the model for 5, 10, and 20 epochs, respectively; after 20 epochs there was no more improvement in accuracy and loss.

**Fig 3.**
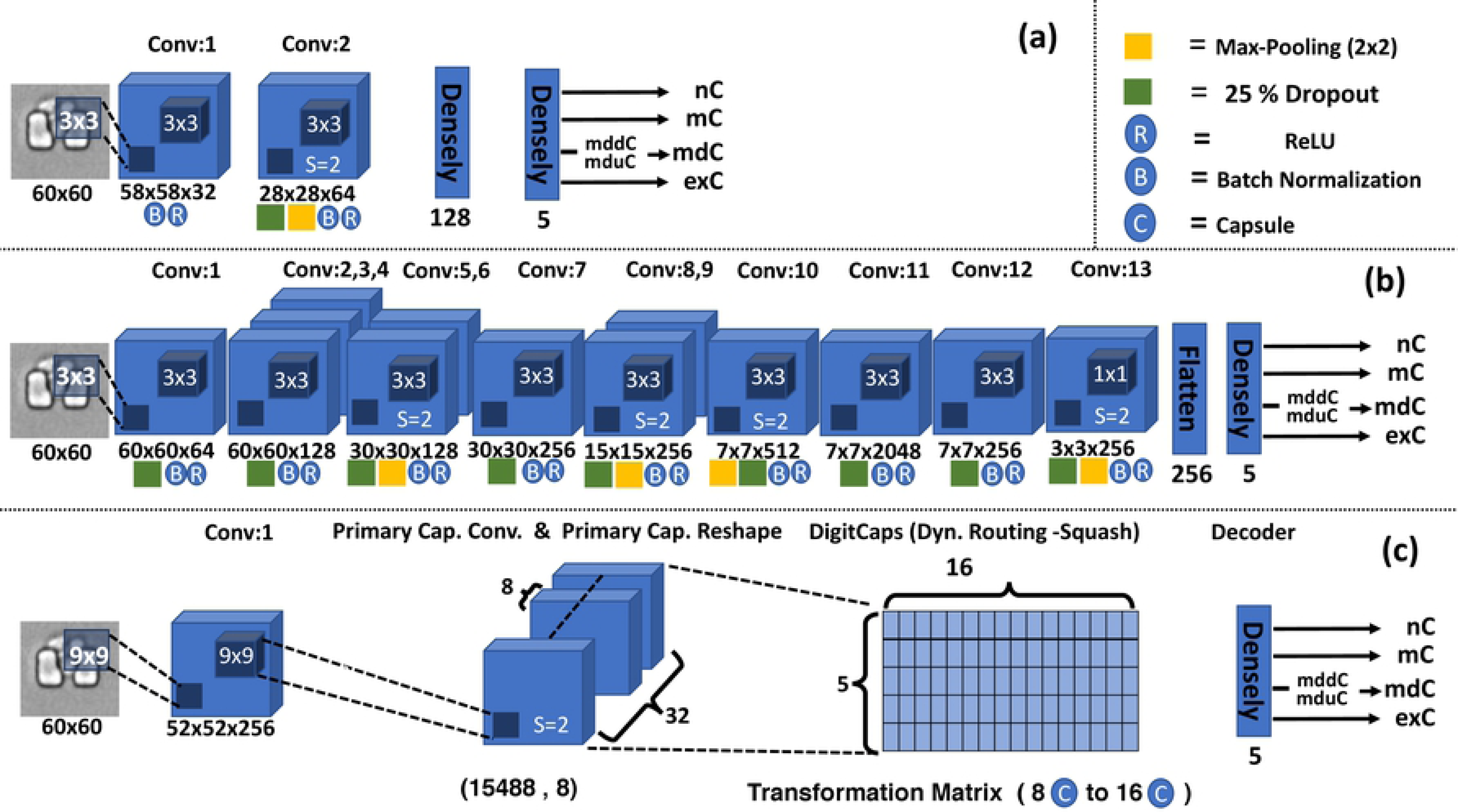
Architectures of three models. (a) CNN-2: A total of 2 convolutional layers and 2 densely connected layers. (b) CNN-13: A total of 13 convolutional layers plus a densely connected layer. (c) Capsule Network: A convolutional layer plus a high-level capsule layer and a densely connected layer. In general, CapsNet contains two parts: the encoder that takes an input image and learns to encode it into 16D instantiating vector parameters, and the decoder that takes a correct DigitCap from a 16D vector and learns to decode it into an original-like image.

### A 13-layered architecture, CNN-13

We are aware of popular examples such as AlexNet [19], VGGNet [20], and GoogleNet [21]. Each of these networks have tens to hundreds of millions of parameters (neural network weights) to learn and require large training datasets. We chose a deep learning architecture, termed the SimpleNet model, as described by HasanPour et al. [22]. HasanPour et al. [22] chose to think of the SimpleNet architecture in groups of layers, where each group of layers is homogeneous and thus can control overall network size and perform specific tasks well, such as classification and object detection. For clarity, we refer to this SimpleNet as CNN-13 in our work. The CNN-13 architecture (see Fig 3 (b)) is a convolutional neural network architecture with 13 layers. CNN-13 has 2–25 times fewer parameters than the popular models. We chose 2×2 and 3×3 kernels for pooling and convolutional layers respectively. We also trained the CNN-13 model for 5, 10, and 20 epochs, and after 20 epochs there was no more improvement in accuracy and loss. In addition, batch normalization and 25% dropout applied to all layers.

### Capsule networks architecture

Capsule networks (CapsNet) is a novel architecture for deep learning. Basic versions of CapsNet have been shown to outperform extremely sophisticated CNN architectures [12]. A previous study showed that CapsNet could classify fluorescent microscopic images [31]. CapsNet replaces the typical pooling layer of CNNs with a more sophisticated weight-routing mechanism. Instead of generating scalar output as used in CNNs, a capsule layer in CapsNet generates a vector as output from convolutional kernel inputs, where the length of the vector represents how likely it is that a feature from the previous layer is present, and the values of the vector are an encoding of all the affine transformation of the kernel inputs. With a more data-efficient architecture (i.e., less information loss), fewer samples are required to train CapsNet models. We used the baseline CapsNet model as in previous works [12, 31] for our comparison studies. Fig 3 (c) shows the architecture of the baseline CapsNet, which contains a convolution layer, primary capsule convolution and primary capsule reshape, DigitCaps (Squash function), and decoder. The kernel size is 9×9 and the stride is 2 for primary capsule convolution. The dimension for primary capsule reshape is 22×22×32 with 8 capsules. A grid search of the hyper-parameters (see S1 Table) led to 108 trained CapsNet models, from which we picked 10 top-performing models. We then examined these 10 models and picked the best-performing CapsNet model for further studies.

### Data augmentation

Due to the tedious process of manual annotation, we have a relatively small number of training images. We tried several affine transformations to augment training images [24–26]. Affine transformations on the original images are a popular and simple augmentation method [26]. The augmentation table for this work is available in S1 Table. We added noise to images and changed brightness, contrast, width, and height of the training images. The total number of trap images in our datasets is 1,000 for each of the five categories. We used 4,104 trap images for training and 896 for testing. We augmented the training images, which resulted in 99,380 training images. The codes and dataset of this work are available from https://github.com/QinLab/GhafariClark2019.

### Performance metrics

Three key metrics have been used in the model analysis [27]. The first is accuracy, e.g., the number of true positive and true negative exC predictions versus all of the exC examples. The second metric is precision, e.g., the true positives prediction of the mC class versus all true positives and false positives of mC. Lastly, we are concerned with a metric called recall [28]. One example of recall is the true positives prediction of the mdC class versus all true positives and false negatives of mdC. Each of these three metrics has its own purpose, and they are oftentimes used together to determine the overall performance of a model [29], written as:

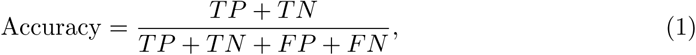

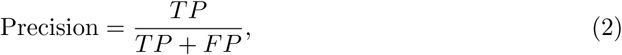

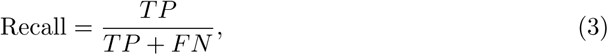

where TP, TN, FP, and FN refer to true positives, true negatives, false positives, and false negatives respectively.

## Results and discussion

### Extension of classes improved the accuracy of predicted biological categories

At the initial stage, we trained and tested all models with four classes: nC, mC, mdC, and exC. Here, mdC refers to any traps with two cells without merging any classes. However, early in the process of model selection and tuning, we discovered that many training images were misclassified when two cells were observed inside the same trap. Hence, some of the best models struggled to reach 60% test accuracy. One approach is to use transfer learning [30] to reduce the misclassifications. Transfer learning is a neural network that starts with pre-trained weights from which models can learn weights in a shorter time. The concept of splitting classes is similar to transfer learning as both methods attempt to make it easy for the models to learn weights; however, these approaches come from different angles. We notice that there are similarities between the exC class and the mdC cell class in cases when the daughter cell is above the mother cell. Based on this observation, we split all images with two cells into two separate classes; in the first class, the daughter cells are on top of mother cells (upward-oriented, mduC class), and in the second class the daughter cells are below the mother cells (downward-oriented, mddC class), as illustrated in Fig 2 (c). At the highest level, creating mddC and mduC classes helped the situation where the neural networks were able to more easily learn the differences of the mduC class and the exC class without having to learn that the mddC and mduC class are the same. It is important to notice that all training and testing activities are based on the computed 5 classes dataset. However, the results for mddC and mduC classes are averaged and labeled as mdC for easier biological understanding as shown in Fig 2 (b).

### CNN-2 performance was improved by training data augumentation

CNN-2 exhibited instability and did not perform well when it was trained with non-augmented training datasets, as seen in blue bars in Fig 4 (a). For example, CNN-2 models trained without augmentation performed poorly on mC, with precision at 71% and recall at 66%. The comparison results in Fig 4 (b) indicate that augmentation mainly improved the accuracy of prediction over the mdC class in this model. As a result of the training data augmentation, the overall accuracy of CNN-2 was improved to 92%. Moreover, the misclassification results show that two common types of misclassifications occurred in CNN-2 while there were only two cells observed inside the trap. For S3 Table CNN-2 (a), the model wrongly predicted two cells instead of three cells due to blurred boundaries. Cases in S3 Table CNN-2 (b, c) were a little more problematic because the CNN-2 model did not recognize the daughter cells above or below the mother cells. Interestingly, for S3 Table CNN-2 (d), the mother cell is almost entirely transparent and ends up not being a problem after recombining the mddC and mduC classes.

**Fig 4.**
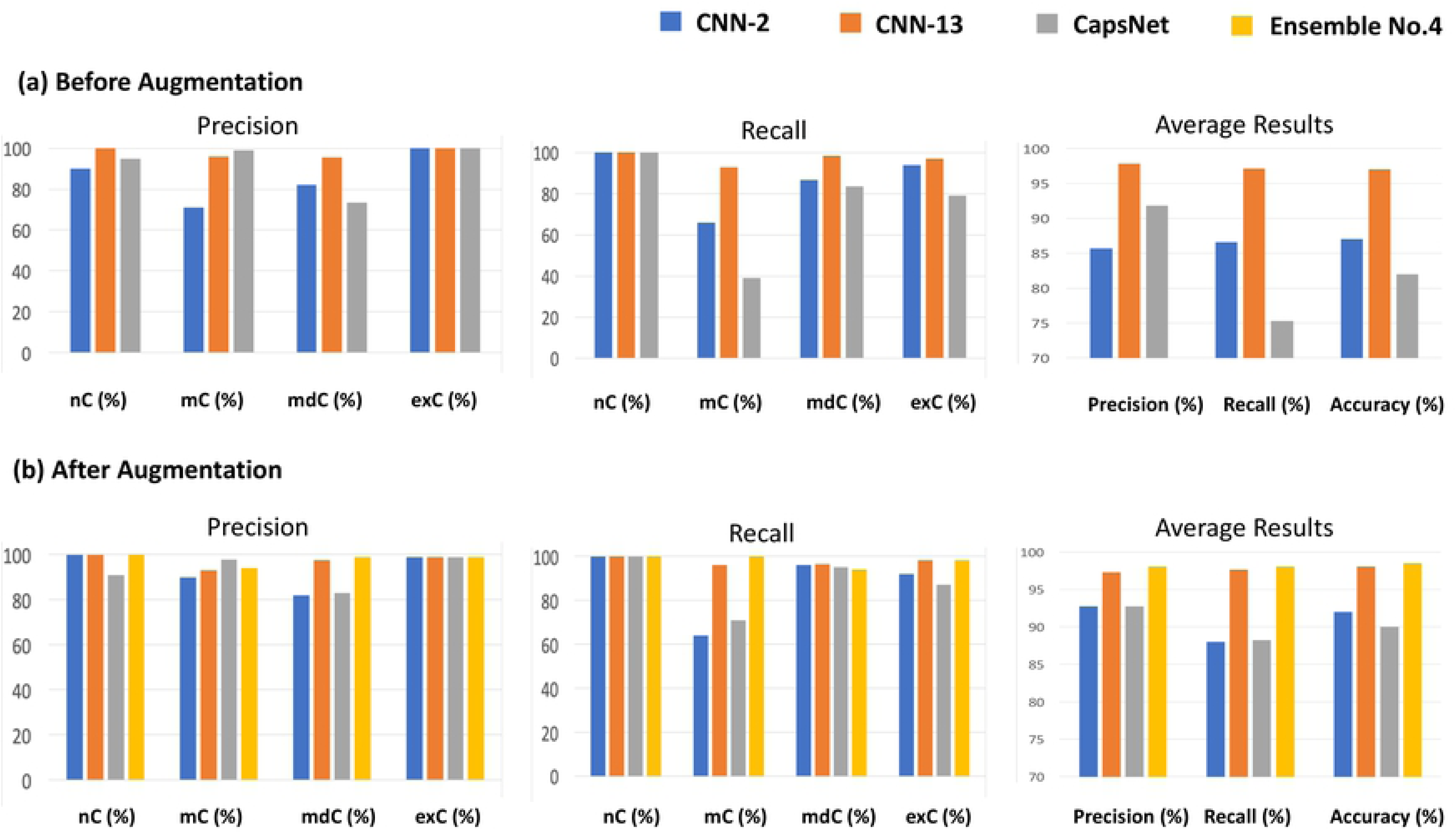
Comparison results for classification models. (a) Combining the mddC and mduC classes increased the performance of the models further. The precision for the mdC class was relatively low, which means that models predicted the mdC class often; however, the prediction was not correct in many of those instances. (b) There are a few elements to note about the model comparison. Every single well-performing model had an augmented dataset. The augmentation mainly improved CNN-2 and CapsNet models.

### CNN-13 performance and impact of training dataset augmentation

CNN-13 showed substantial improvement in average accuracy in comparison to CNN-2, and this improvement occurred for CNN-13 models trained with and without augmentation of training datasets, as shown in Fig 4. Augmentation of training data also led to more stable CNN-13 models as seen when changes of the cost functions during training became more smooth with augmented datasets. Surprisingly, augmentation had a marginal effect on the accuracy, precision, and recall of CNN-13. Before augmentation, the model predicted 100% on nC class (precision and recall) and exC class (precision). Most of the misclassification appears to be in the mC and mdC classes. After augmentation, prediction for nC did not change (100%) and mC recall improved from 93% to 96% (precision had the opposite reaction). Furthermore, S2 Table shows that augmentation had a slight improvement in the mC and exC classes but a negative effect on the mdC class. The overall accuracy for this model was 97% (before augmentation) and 98% (after augmentation) respectively as shown in orange bars.

Considering misclassification for CNN-13, S3 Table CNN-13 (a) shows several cells clustered together. After further inspection, this image was classified with near 100 % certainty. Although this instance is uncommon, it still poses problems in cell type identification. The mistake on S3 Table CNN-13 (b) is more understandable because there is a mother cell with seemingly two daughter cells on top. The algorithm did not classify this example in the exC class and instead predicted it as mduC. Since one of these cells could actually be a true daughter cell, this image may not be as problematic. Image S3 Table CNN-13 (c) is similar to the previous image, but the boundary between the two cells on top of the mother cell are so thin that it is reasonable to think that it is a deformed single daughter cell to the untrained eye. Finally, S3 Table CNN-13 (d) illustrates a mistake that was common in the CNN-2 model where mduC or mddC were predicted as mC due to blurred boundaries.

### CapsNet performance and impact of training datset augmentation

We found that the performance of CapsNet was more sensitive to hyper-parameters than were the CNN-2 and CNN-13 models, based grid searches on the hyper-parameters detailed in S1 Table. We picked the best-performing CapsNet model for this study. The training dataset augmentation mainly improved CapsNet’s accuracy of the mC category but not in other categories, as shown in Fig 4. The overall accuracy of CapsNet reached 90% after augmentation. In Zhang et al. [31], a close range of accuracy was reported for fluorescent images.

In case of misclassification, S3 Table CapsNet (a) shows that there is a small cell on the top right portion of the mother cell that seemed to be overlooked by the CapsNet model. One potential cause for this misclassification is that the two cells on top of the mother cell are quite different in size. S3 Table CapsNet (b) is one of the problematic misclassifications that CNN-13 was good at detecting. S3 Table CapsNet (c) shows a transparent cell that could be dead or senescent. This type of image is unlikely to happen often enough for the model to learn effectively. S3 Table CapsNet (d) shows another interesting example. It looks as though a mother cell was too big for the trap and is reproducing daughters that flow over the outside edge of the trap.

### Deeper layers bring moderate improvement and challenging performance of the CapsNet

Surprisingly, CNN-2 can predict the nC category with 100% accuracy, even though it has a skeleton architecture (see the confusion matrix S3 Fig). We found that performance of CNN-2 can be greatly improved by augmentation and adding layers. As expected by the increased number of layers, CNN-13 had greater overall accuracy than CNN-2, as shown by its confusion matrix (see S3 Fig). With the additional 11 more layers and much more training time, CNN-13 improved the overall accuracy to 98%, a moderate 6% increase from CNN-2. We also found that performance of CNN-13 is not substantially changed by applying augmentation. Fig 5 shows that augmentation improved the total prediction by 0.22%, which is around 16 times lower than CNN-2 and decreased the total mis-prediction by 8.3%, which is considerably lower than CNN-2. On the other hand, CapsNet was the weakest model in terms of average accuracy. According to the confusion matrix (see S3 Fig), the model had only great prediction for nC (180/180). Surprisingly, the model had the best prediction (354/360) for the mdC class before any augmentation where both CNN-2 and CNN-13 struggled with prediction (with or without augmentation). However, the model had poor prediction for the mC and exC classes. Fig 5 illustrates that the augmentation was an effective approach that improved the total prediction by 7%, which is almost double CNN-2 model and decreased the total mis-prediction by 30.8%, better than the other two models. In other words, CapsNet is much more sensitive to data augmentation than the other two CNN models are, and it can preform well on a specific class.

**Fig 5.**
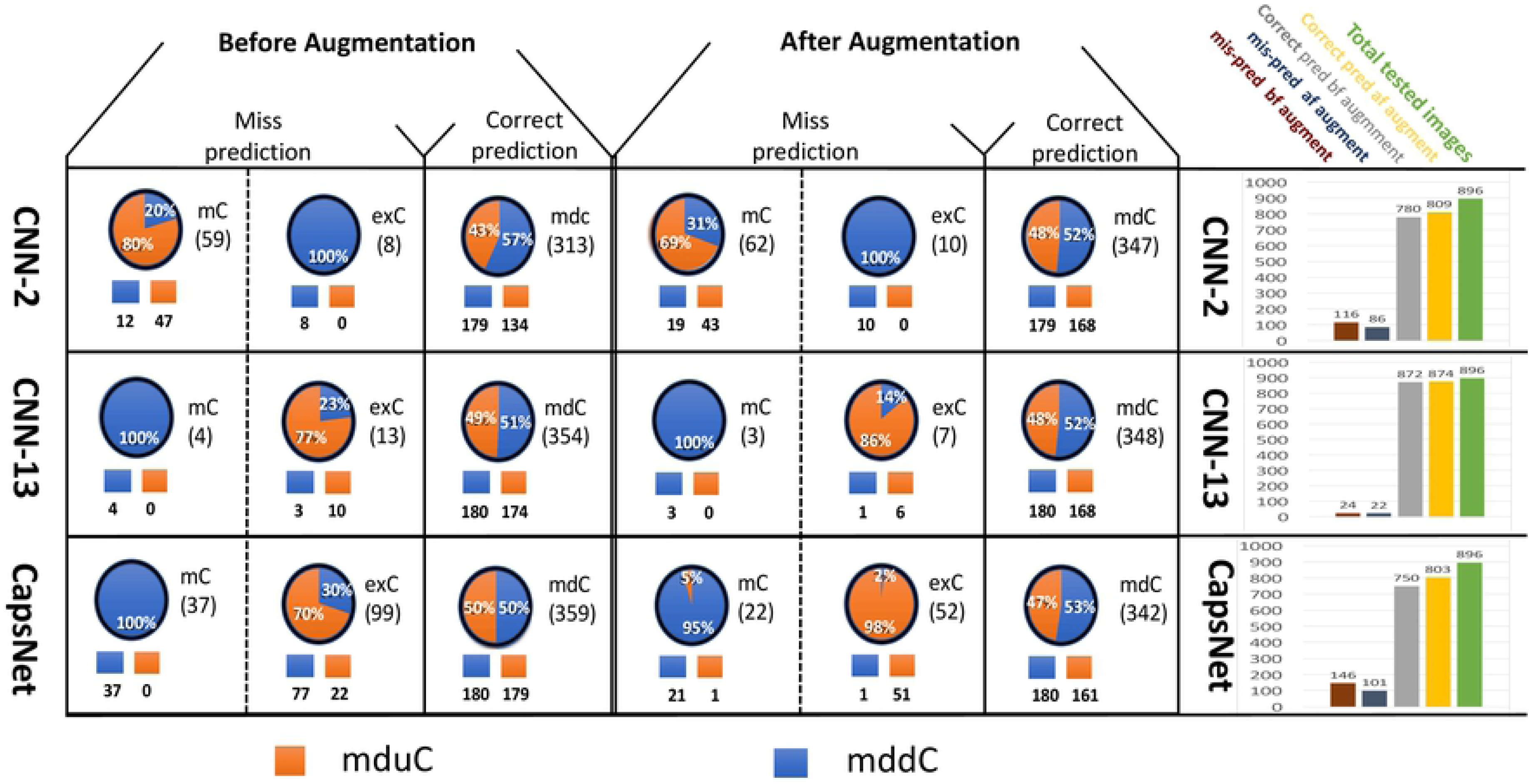
Each of the three deep learning models has idiosyncratic error profiles. Table presenting the example of predicted mdC class that included correct predictions (mdC) and mis-predictions (mC and exC). Based on the mdC columns in the confusion matrix (S3 Fig), predicted mdC are broken down to mddC and mduC extended classes. Orange areas indicate predicted mduC, and blue areas indicate predicted mddC. In the bar graphs, total tested images are in green. Correct predictions of all 4 biological classes after augmentation are in yellow. Correct predictions of all 4 biological classes before augmentation are in gray. Mis-predicted classes after augmentation are in blue. Mis-predicted classes before augmentation are in red. These results motivated us to generate ensemble models.

### Each deep learning model has its own profiles of misclassifications

We also investigated misclassification behavior of individual models for the mC, mdC, and exC classes (see Fig 5). In terms of correct-prediction balance between mddC and mduC, Fig 4 demonstrates that all the models had close balance of prediction for mddC and mduC (before and after augmentation). In terms of mis-prediction, the CNN-2 model had opposite behavior of the CNN-13 and CapsNet models. For CNN-2, the mduC class had a higher percentage of misclassification for the mC class, and the exC class had higher misclassification for the mddC class. For CNN-13 and CapsNet, the mddC class had a higher percentage of misclassification for the mC class, and the exC class had a higher misclassification for mduC. These comparisons indicate why we consider an ensemble model as an alternative. For Ensemble No. 4, each of the misclassifications of the ensemble are obviously misclassifications of at least one of the three models.

### Ensemble models performance

Because each single deep learning model had uneven performance in the 4 biological categories, we thought combining them may lead to better performance. There are four different ways to combine the three single deep learning models (see S2 Fig). We chose a straightforward ensemble method to weight the predictions of each model based on their overall accuracy [32] and misclassifications. Specifically, CNN-13 has the highest prediction weight, followed by CNN-2 and then CapsNet. We found the three-member ensemble, No. 4, outperformed all of the two-member ensembles. The overall accuracy of ensemble No. 4 is 98.5% as shown in Fig 4 (b), in yellow.

## Future work

While correctly classifying images into one of the four discussed categories was the focus of this work, there are still improvements to be made such as the image pre-processing differently from data augmentation. In addition, we could improve the overall ensemble by adding more diversity to the set of models. For example, the sequential nature of the problem could lend itself nicely to a Last Short-Term Memory (LSTM) architecture [33].

## Conclusion

We compared three deep learning models for classification of microfluidic images of dividing yeast cells. Microfluidiic images are typically low resolution, which poses challenges for computational analysis. We found that augmentation of training data can improve performance of both convolutional and capsule networks. We found that extended computed classes could improve performance of deep learning methods for classifying biological classes. We found that a baseline architecture of convolutional network with two layers could give 92% overall accuracy. We found that deep layered convolutional networks could improve the overall accuracy at the expense of substantially more computing cost. We found that a baseline architecture of capsule neural networks did not outperform the deep-layered convolutional networks in terms of overall accuracy, though the baseline capsule networks could detect a specific type of data with better performance. Consequently, an ensemble model reached 98.5% overall accuracy by combining the strengths of different models. Overall, we found that convolutional and capsule neural networks have complementary performance for classification of microfluidic images of dividing yeast cells.

## Acknowledgments

This work is partially supported by the NSF CAREER award #1453078 (transferred to #1720215), NSF award #1761839, a departmental start-up fund, internal CEACSE awards from the University of Tennessee at Chattanooga, and the computing facility of the SimCenter at the University of Tennessee at Chattanooga. We also acknowledge the support of NIH grants #R01AG052507 and #R42AG058368.

## Supporting information

**S1 Fig. Microfluidics Images.** (a) Time-lapsed images from time-point 001 to time-point 391. Black circles with connected dash-lines indicate that some of traps become overcrowded over time. (b) Each imaged partitioned to 60ox60 pixels sub-image and individual trap image is highly variable. While traps and cells have a limited number of orientations, the contrast, brightness and image quality all add great complexity to the dataset. There are often shadows, depending on the lighting conditions of the experiment.

**S2 Fig. Ensemble Models Combination.** Results of CNN-2, CNN-13 and CapsNet models indicated that there are numerous ways to ensemble (i.e. combine) models together in order to create a single aggregate model. We explored the results from all possible ensembles with different combinations based on practical and key performance metrics.

**S3 Fig. Models confusion matrix.** Three models confusion matrix with indication of augmentation effectiveness.

**S1 Table. Grid search and augmentation options.** A total of 108 combinations were trained without and with augmented datasets.

**S2 Table. Models comparison for performance Metrics.** The results of precision, recall and accuracy for all models.

**S3 Table. Sample image of most common misclassifications.**

CNN-2 (a) label exC: prediction mduC, CNN-2 (b) label mduC: prediction mC, CNN-2 (c) label mddC: prediction mC, CNN-2 (d) label mduC: predicted mddC.

CNN-13 (a) label exC: prediction mduC, CNN-13 (b) label exC: prediction mduC, CNN-13 (c) label exC: prediction mduC, CNN-13 (d) label mddC: prediction mC.

CapsNet (a) label exC: prediction mduC, CapsNet (b) label mddC: prediction mC, CapsNet (c) label mC: prediction mduC, CapsNet (d) label mduC: prediction exC.

Ensemble No.4 (a) label mdC: prediction mC, Ensemble No.4 (b) label exC: prediction mdC, Ensemble No.4 (c) label exC: prediction mdC, Ensemble No.4 (d) label mdC: prediction exC.

